# Expanding the toolbox of broad host-range transcriptional terminators for Proteobacteria through metagenomics

**DOI:** 10.1101/485938

**Authors:** Vanesa Amarelle, Ananda Sanches-Medeiros, Rafael Silva-Rocha, María-Eugenia Guazzaroni

**Affiliations:** Department of Microbial Biochemistry and Genomics, Biological Research Institute Clemente Estable, Montevideo, Uruguay; FMRP - University of São Paulo, Ribeirão Preto, SP, Brazil; FFCLRP - University of São Paulo, Ribeirão Preto, SP, Brazil

**Keywords:** transcriptional terminators, functional metagenomics, synthetic biology, biological parts, Proteobacteria

## Abstract

As the field of synthetic biology moves towards the utilization of novel bacterial chassis, there is a growing need for biological parts with enhanced performance in a wide number of hosts. Is not unusual that biological parts (such as promoters and terminators), initially characterized in the model bacteria *Escherichia coli*, do not perform well when implemented in alternative hosts, such as *Pseudomonas*, therefore limiting the construction of synthetic circuits in industrially relevant bacteria. In order to address this limitation, we present here the mining of transcriptional terminators through functional metagenomics to identify novel parts with broad host-range activity. Using a GFP-based terminator trap strategy and a broad host-range plasmid, we identified 20 clones with potential terminator activity in *Pseudomonas putida*. Further characterization allowed the identification of 4 unique sequences between 58 bp and 181 bp long that efficiently terminates transcription in *P. putida, E. coli, Burkholderia phymatum* and two *Pseudomonas* strains isolated from Antarctica. Therefore, this work presents a new set of biological parts useful for the engineering of synthetic circuits in Proteobacteria.

## Introduction

Bacteria play a central role in the biotechnology industry and the field of synthetic biology is rapidly growing towards the generation of novel tools and protocols for the engineering of these organisms ^1,2^. In order to take advantage of their full potential, it is essential to have an efficient set of tools that can be used in organisms with relevant features related to the application intended. For example, *E. coli* is a classical host in terms of protein production and metabolic engineering ^3^, while gram-positive bacteria such as *Streptomyces* hold the potential for the production of small bioactive molecules ^4^. By the same token, the natural features of *Salmonella* make them a suitable host for tumor targeting applications ^5^, while the metabolic robustness and versatility of *Pseudomonas* make these bacteria promising chassis for harsh and intense industrial conditions ^6^. Yet, most of genetic tools have been initially constructed for *E. coli* and it is not unusual that biological parts required for synthetic circuit assemble are not fully functional in alternative hosts ^2^. In these cases, the initial failure of synthetic circuits constructed in alternative hosts leads to the need of intense re-adaptation of these parts to the new hosts ^7,8^. Therefore, as synthetic biologists turn towards non-classical bacterial chassis, there is a growing demand for orthogonal systems that could efficiently operate in a broad number of hosts ^9,10^. Attempts in this direction have been made towards the construction of novel expression systems ^11^, transformation/DNA delivery strategies ^12,13^ and plasmid vectors ^14^, among others.

Recently, several reports have described the construction of novel expression systems, based on inducible as well as constitutive promoters, both with narrow or wide host-range ^7,10,15,16,2^. Yet, while much is known about transcription initiation, considerably less information is available about transcriptional termination ^17,18^. In Bacteria, transcription termination occurs mainly through two mechanisms. In intrinsic termination – also termed Rho-independent termination –, strong secondary structures are formed at the 3’ end of nascent RNA transcript which lead to the detachment of the RNA Polymerase (RNAP), and this mechanism does not seem to require any additional *trans* acting element ^17^. On the other hand, Rho-dependent termination requires a helicase which navigates the mRNA and actively detaches paused RNAPs ^19^. Similar mechanisms are found in Archaea and Eukarya ^20^. For the construction of complex synthetic circuits, efficient transcription termination sequences are require to ensure insulation of each part of the system. Failure in this process could lead to unwanted regulatory interactions. Yet, while some recent works have systematically characterized intrinsic terminators in bacteria, this has been mainly performed in the model organism *E. coli* ^21,22^. Therefore, there are few well-characterized fully functional terminator sequences that can be used in different bacterial hosts. In this sense, functional metagenomics holds the potential to access a great diversity of biological parts that can be used in a number of applications ^23,24^. In this work, we search for novel transcriptional terminators in soil microbial communities by a functional metagenomic approach. Using a GFP-based terminator trap broad host-range vector, we identified several candidate sequences by the functional screening of a metagenomic library constructed in *Pseudomonas putida*. We demonstrated that these new terminator sequences are functional in several Proteobacteria, including *E. coli*, and this new set of tools could be useful for novel synthetic biology projects in non-classical chassis.

## Results and discussion

### Mining transcriptional terminators in metagenomic libraries

As presented above, while many transcriptional terminators have been well-characterized in *E. coli*, only few functional terminators are available for non-classical bacteria. As an example, the widely used minimal lambda T1 terminator from *E. coli* is not fully functional in *P. putida* KT2440, as this element allows readthrough when placed in broad host-range vectors (**Fig. S1**). In this sense, we hypothesize that an optimal terminator screening strategy would require these elements to be first validated in the organism of interest and only then assayed in *E. coli*. The overall strategy used in this work is schematically represented in **Fig. 1**. In order to identify novel transcriptional terminator elements, we constructed a metagenomic library from eDNA extracted from soil samples. For this purpose, a strong constitutive promoter (BBa_J23100) controlling the expression of a GFPlva reporter was cloned into the broad-host range vector pSEVA231 ^14^. The resulting reporter vector was used for terminator trapping by cloning the eDNA in the region between the promoter and the ribosome binding site of GFPlva **(Fig. 1)**. The constructs were introduced in *P. putida* KT2440 and the metagenomic library was screened for the lack or reduction in GFP signal. Using this approach, a total of 163 colonies were screened and 20 colonies with apparent GFP signal reduction/absence were selected for further validation. Plasmid DNA extraction, digestion and DNA sequencing revealed that all 20 clones presented a DNA fragment ranging from 58 to 424 bp, with an average insert length of 211 bp (**Table S1**). These clones were randomly named T1 to T20 and were used for further experimental validation as described below. **Table S2** provides information about sequence similarity, both at nucleotide and amino-acid level, of the metagenomic inserts.

**Figure 1.**
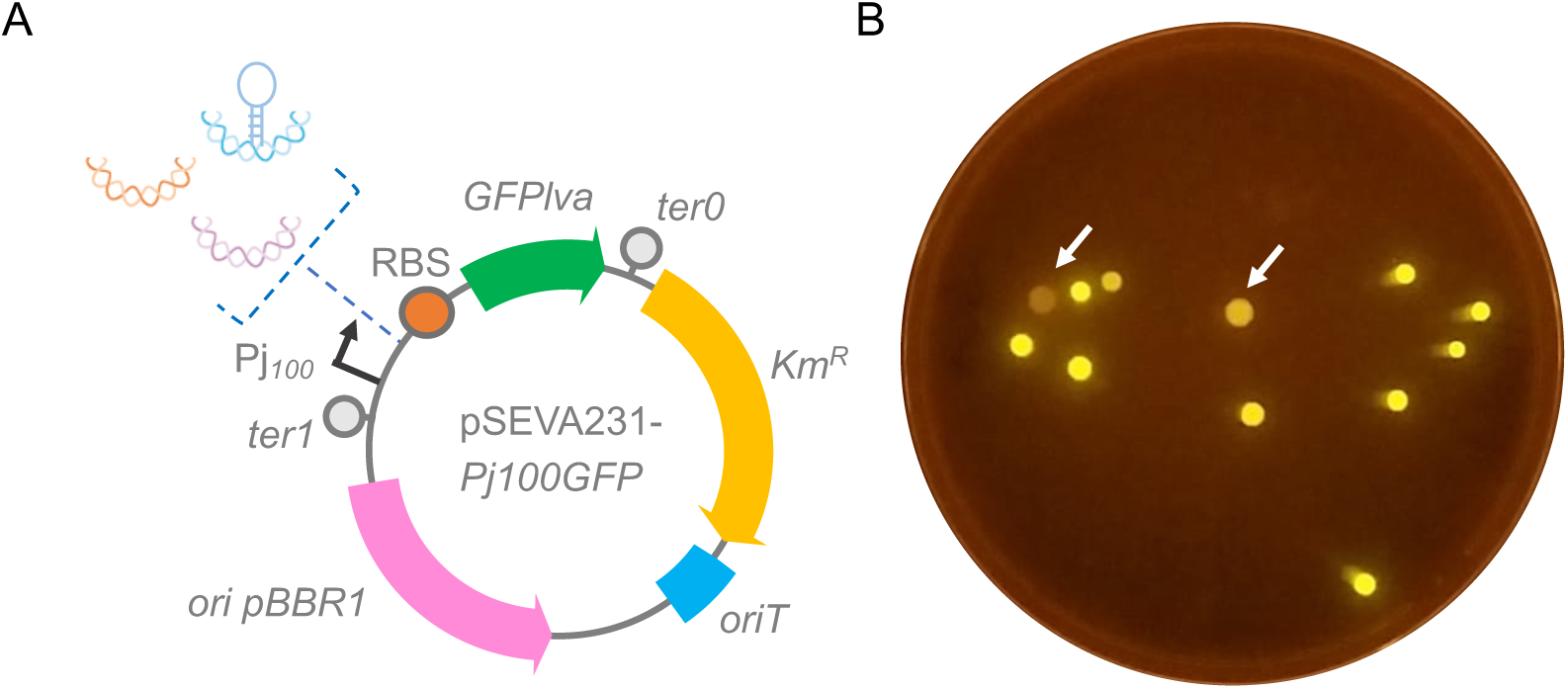
Schematic representation of the strategy used in this study. **A)** Metagenomic DNA fragments carrying potential transcriptional terminators were cloned between the strong promoter P*j_100_* and the RBS (ribosome-binding site) in the reporter trap-vector pSEVA231-*Pj100*GFP. Functional elements of the plasmid backbone are shown: Km^R^, antibiotic resistance marker; *oriT*, origin of transfer; *ori* pBBR1, broad host-range origin of replication; *ter1* and *ter0*, transcriptional terminators. **B)** Metagenomic library clones presenting lack or reduction in GFP signal, in a blue-light transilluminator, are indicated by arrows.

Diverse algorithms (Mfold and ARNold, FindTerm ^25^) were applied in order to identify potential intrinsic terminators present in all the eDNA sequences cloned. FindTerm retrieved T18 as the only putative Rho-independent terminator, while the analysis of RNA secondary structure with Mfold allowed identifying a putative typical Rho-independent terminator in T1 sequence. As these algorithms are designed to search for typical Rho-independent transcriptional terminator motifs, a hairpin structure followed by a poly-U tract, the presence of Rho-dependent terminators or even unusual Rho-independent terminators in the metagenomic sequences cannot be discarded, considering the diverse phylogenetic origin of the DNA sequences screened ^26,27^.

### The novel transcriptional terminators display broad host-range activity

Once we identified candidate terminators in *P. putida*, we further characterize these sequences by assessing *in vivo* terminator activity in liquid media. As shown in **Fig. 2A**, for seven selected candidate terminators all strains presented reduced GFP expression when compared to the positive control harboring pSEVA231-*Pj100*GFP plasmid, where no sequence is inserted between the strong promoter and the reporter gene. While some sequences presented only partial terminator activity (T2 and T3), others presented expression levels very close to the negative control harboring pSEVA231 plasmid (such as for T1, T7 and T9). It is worth noting that these three sequences were very short (between 58 and 181 bp long) (**Table S1**), resulting thus into promising new sequences for transcriptional termination in *P. putida*. The terminator effect was maintained for all constructs over a period of 8 hours (**Fig. 2B**). As shown in **Fig. 2C**, the presence of eDNA in the constructs has no significant effect in the growth pattern when compared to the presence of pSEVA231-*Pj100*GFP and pSEVA231 plasmids. The characterization of the remaining 13 candidate sequences evidenced a wide range of termination efficiencies (**Fig. S2**).

**Figure 2.**
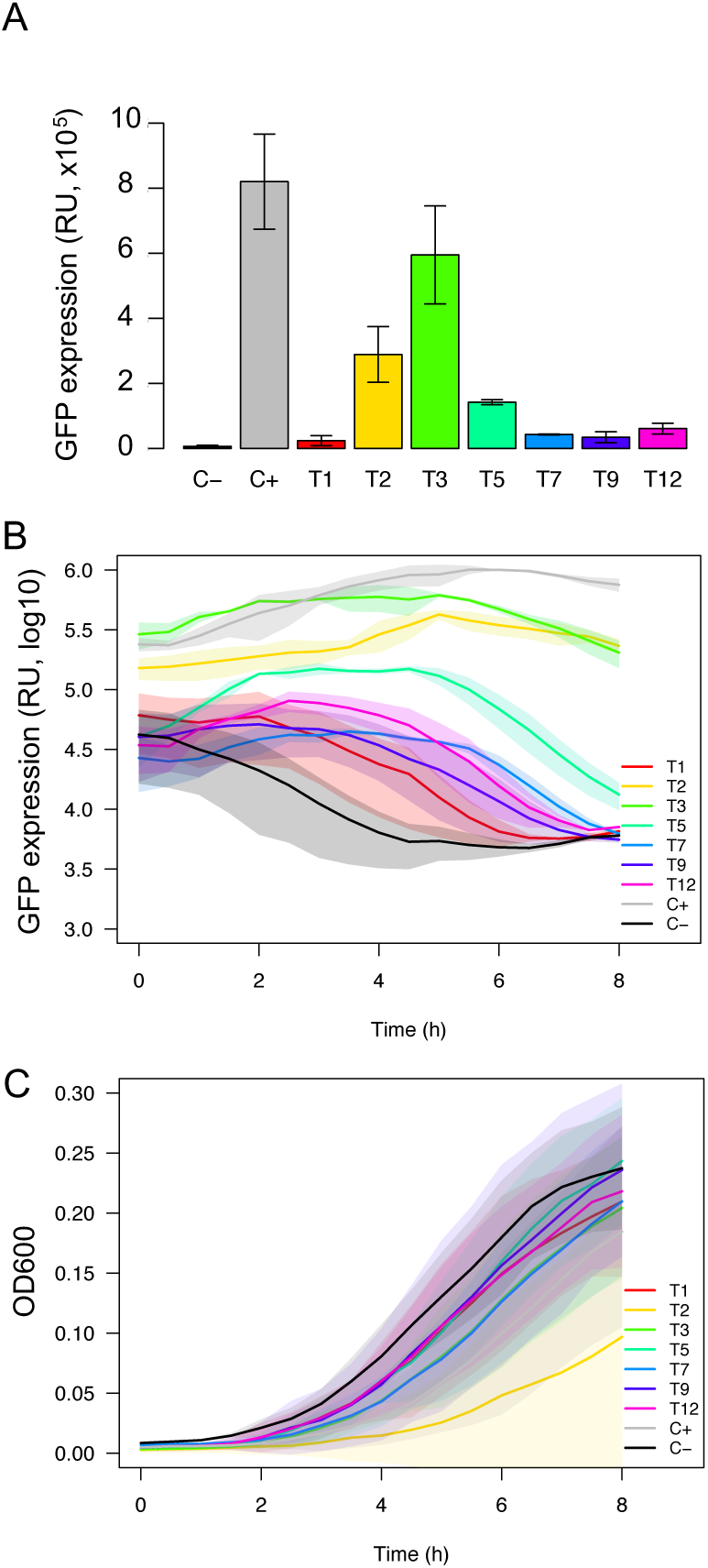
Characterization of novel terminators in *P. putida* KT2440. In order to characterize the transcriptional termination effect in selected metagenomic clones, *P. putida* KT2440 strains harboring the different reporter plasmids were grown in liquid M9-glymedia during 8h and the optical density at 600 nm (OD600) and fluorescence (GFP) were measured every 30 min. Strains harboring either pSEVA231 or pSEVA231-Pj100GFP were used as positive (C+) and negative (C−) controls, respectively. **A)** Terminator activity at 4h **B)** Terminator activity (in log10 scale) over time. **C)** Growth curve of the strains. All graphs represent the average from three independent experiments. Standard deviation from three experiments is represented as vertical bars in A and as shaded regions in B and C.

Once we analyzed the terminators in *P. putida*, we investigated their functionality in four additional Proteobacteria host. For this purpose, we introduced the plasmids presented in Fig. 2 into *E. coli* DH10B, *B. phymatum* STM815^T^ and two psychrotolerant *Pseudomonas* strains isolated from Antarctica ^28^, named as *Pseudomonas spp. UYIF39* and *Pseudomonas spp*.

*UYIF41*. As shown in **Fig. 3A-B**, all tested *E. coli* strains presented GFP expression levels virtually undistinguishable from the negative control, indicating that all 7 terminator sequences were fully functional in this host. This result strength the notion that functional terminator in *E. coli* are not always active in other hosts such as *P. putida*. It is worth mentioning that plasmids were extracted from *E. coli* and the presence of the terminator sequences originally identified in *P. putida* were confirmed by PCR. When these terminators were analyzed in *B. phymatum* and the two psychrotolerant *Pseudomonas* strains (**Fig. 3C-H**), the termination profile was very similar to that of *P. putida*, where T1 was the most effective terminator in all strains and a host-dependent efficiency for T7, T9 and T12 was observed. Also, the introduction of these constructs in all four strains did not cause any defect in growth under the experimental conditions assayed (**Fig. S3**). Taken together, these results demonstrate that the novel terminators identified in this work are highly functional in a broad range of bacterial strains.

**Figure 3.**
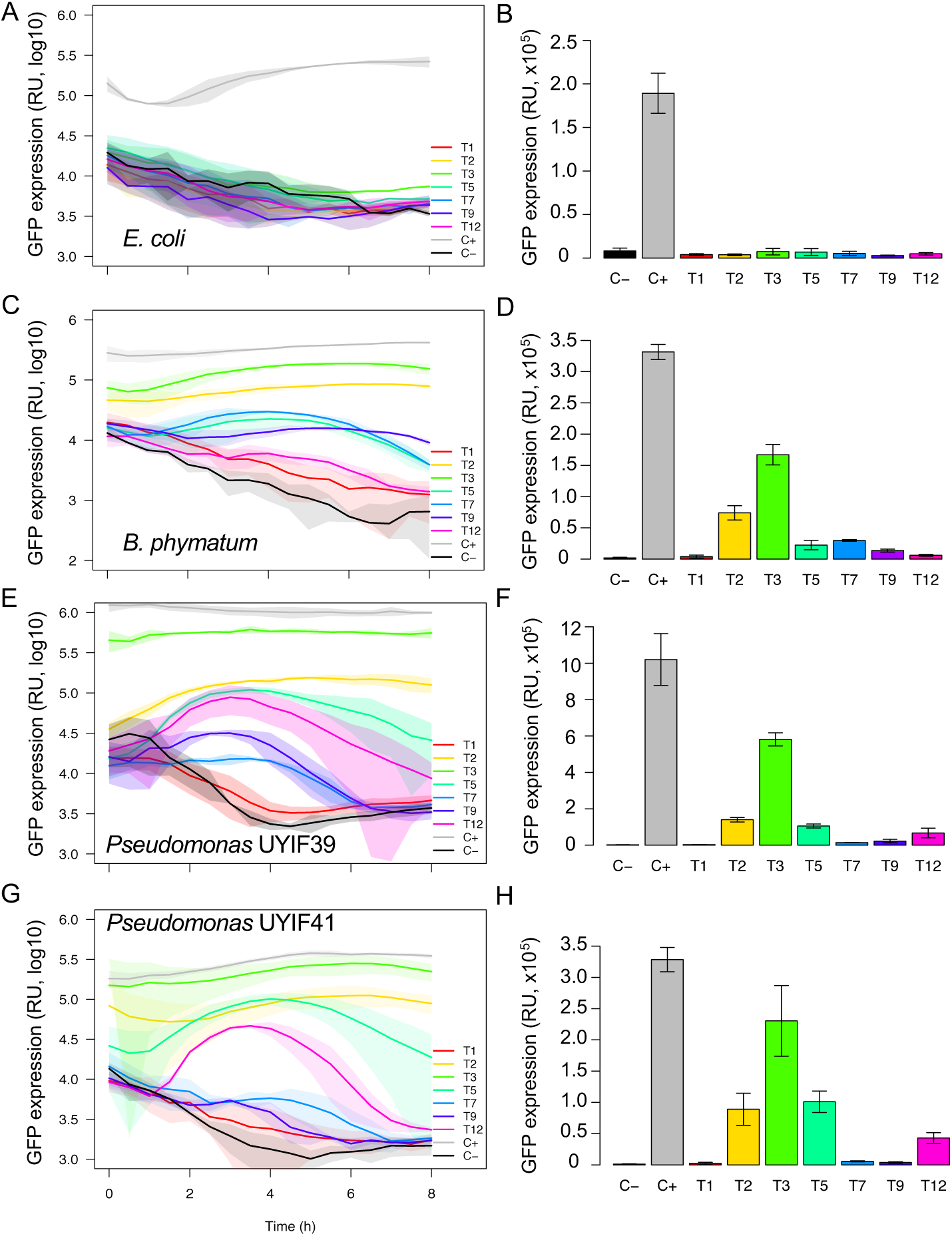
Characterization of novel terminators in alternative hosts. In order to characterize the transcriptional termination effect in alternative hosts other than *P. putida*, plasmids were transferred to *E. coli, B. phymatum* and the psychrotolerant *Pseudomonas* UYIF39 and UYIF41. All the strains were grown in liquid M9, with the appropriate carbon source, during 8h and the optical density at 600 nm (OD600) and fluorescence (GFP) were measured every 30 min. Strains harboring either pSEVA231 or pSEVA231-Pj100GFP were used as positive (C+) and negative (C−) controls, respectively. **A)** Terminator activity (in log10 scale) of *E. coli* strains over time. **B)** Terminator activity of the *E. coli* strains at 4h. **C)** Terminator activity (in log10 scale) of *B. phymatum* strains over time. **D)** Terminator activity of *B. phymatum* strains at 4h. **E)** Terminator activity (in log10 scale) of the *Pseudomonas sp*. UYIF39 strains over time. **F)** Terminator activity of the *Pseudomonas sp*. UYIF39 strains at four 4h. **G)** Terminator activity (in log10 scale) of the *Pseudomonas sp*. UYIF41 strains over time. **H)** Terminator activity of the *Pseudomonas sp*. UYIF41 strains at 4h. All graphs represent the average from three independent experiments. Standard deviation from three experiments is represented as shaded regions in A, C, E and G, and as vertical bars in B, D, F and H.

### The novel terminators are functional at single cell level

While most characterization of biological parts are usually performed at population levels, the role of cell individuality is widely recognized as crucial for network performance ^29^. In this sense, bacterial expression systems are usually categorized as having graded or all-or-none behavior, and these have strong consequences for the final circuit dynamics ^30,31^. Therefore, we decided to investigate how the newly-identified terminators behave at the single-cell level. For this purpose, the three best terminators (T1, T7 and T9) were assayed by flow cytometry in the five hosts. As shown in **Fig. 4**, while terminator T1 displayed a low-expression dynamics with a single population of cells similar to the negative control in all strains, terminators T7 and T9 displayed a low-expression population with broader distribution than the negative control, being distribution strain-dependent. In this sense, strains *P. putida*, *B. phymatum* and *Pseudomonas* spp.UYIF39 were more heterogeneous, yet with a single population present. Taken together, these results demonstrated that three of the strongest terminators identified in this work are efficient in transcription termination at the single-cell level as none of them displayed both, active and inactive, populations.

**Figure 4.**
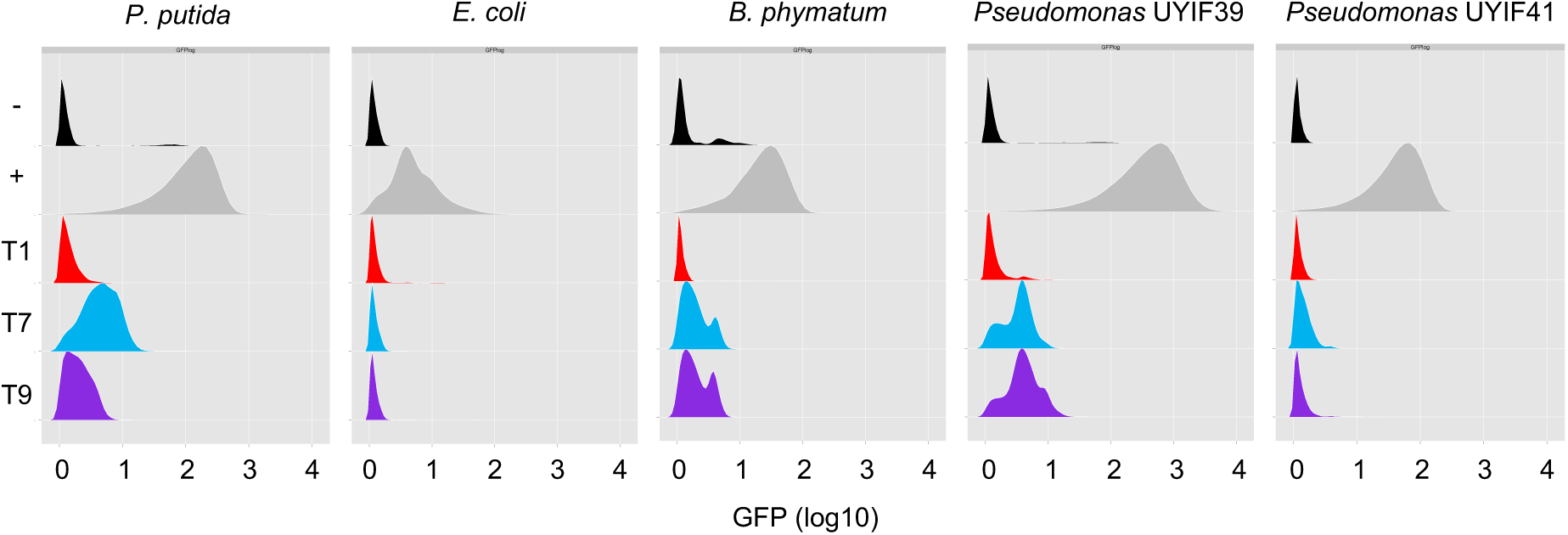
Flow cytometry analysis of three broad-host range terminators in different hosts. Strains were gown in M9 media supplemented with an appropriate carbon source for 4 hours and analyzed by flow cytometry. Data from 20,000 events were collected and processed using flowCore and flowVIz packages in R. The results are representative of three independent experiments. Strains harboring either pSEVA231 or pSEVA231-Pj100GFP were used as positive (+) and negative (-) controls, respectively.

## Conclusions

In this work, we present a strategy for the mining of novel functional terminator sequences in Proteobacteria other than *E. coli*. Using a functional metagenomic approach, twenty terminators with different efficiencies and host-dependency were identified, being the most promising ones less than 200 bp. This set of new, short, and wide host-range terminator sequences could be crucial for future synthetic biology projects requiring complex circuits, since re-use of parts could lead to DNA rearrangements due to homologous recombination events. Also, the fact that these new elements are functional in several bacteria allows the user to construct and test the circuit in one host and then implement it in a different one without the loss of functionality. We anticipate that similar functional metagenomic approaches could be used in the future to mine other broad host-range elements, such as RBS and promoters, increasing the toolbox for the construction of truly orthogonal synthetic circuits.

## Methods

### Bacterial strains, plasmids and growth conditions

Bacteria and plasmids used in this study are detailed in supplementary **Table S3**. *E. coli* and *Pseudomonas* strains were grown aerobically at 30 °C and 200 rpm in either LB or M9 minimal media ^32^ supplemented with 0.1 mM casamino acids and 1% glycerol as carbon source (M9-gly). *Burkhordelia* strains were grown in either TY ^32^ or M9 minimal media supplemented with 0.1 mM casamino acids and 0.2% citrate. When required, 50 μg/ml kanamycin (Km) or 25 μg/ml chloramphenicol (Cm) were added to the media.

### Construction of the metagenomic library

In order to build the terminator trap vector, a 857 pb DNA fragment harboring the GFPlva protein under the control of a strong synthetic constitutive promoter (BBa_J23100) was obtained from pMR1-*Pj100* plasmid ^33^, after digestion with HindIII/EcoRI. The resulting *Pj100GFP* region was later cloned at the HindIII/EcoRI sites of pSEVA231 ^14^, generating plasmid pSEVA231-*Pj100*GFP that was used for terminator trapping in environmental metagenomic DNA (eDNA). For this purpose, previously isolated eDNAs ^34^ obtained from soil samples were partially digested with Sau3AI enzyme to produce fragments between 100 bp and 400 bp that were cloned at the BamHI site of pSEVA231-*Pj100*GFP, which is located between *Pj100* promoter and the ribosome binding site (RBS) of GFPlva gene. The constructs were introduced in *P. putida* by electroporation. The metagenomic library was plated in LB Km and colonies were screened for the lack or reduction in GFP signal in a blue-light transilluminator. Colonies harboring putative terminator sequences from eDNA origin were selected for further analysis.

### Analysis of transcriptional terminator activity

In order to quantify the termination efficiency, *P. putida* harboring putative transcriptional terminators were used to asses GFP expression *in vitro*. Single colonies were grown overnight in M9-gly at 30°C and 200 rpm. The overnight growth was placed in a 96-well plate diluted for 1:20 in 200 μl of M9-gly. The analysis was made using a Victor X3 plate reader (PerkinElmer) over a period of eight hours at 30 °C, measuring every 30 minutes the optical density 600 nm (OD) and fluorescence (excitation 488 nm and emission 535 nm). *P. putida* harboring either pSEVA231 or pSEVA231-Pj100GFP were used as negative and positive controls, respectively. All assays were done in technical triplicates in the same plate and in biological triplicates in different experiments.

To assess the effectiveness of the terminator sequences found in *P. putida* in other Proteobacteria hosts, seven constructs were selected based on the strength of the putative terminator effect. Five plasmids harboring efficient terminator sequences (T1, T5, T7, T9, T12) and two plasmids harboring medium terminators (T2 and T3) were obtain from *P. putida* and used to transform *E. coli, Pseudomonas* UYIF39, *Pseudomonas* UYIF41 and *B. phymatum* by electroporation. As negative and positive controls, all strains were transformed with either pSEVA231 or pSEVA231-*Pj100*GFP, respectively. For quantification of the termination efficiency in these hosts, GFP expression was assessed *in vitro* as mentioned before for *P. putida*, performing it at 30 °C for *E. coli*, 30 °C for *B. phymatum* and 25 °C for *Pseudomonas* UYIF39 and UYIF41. For *B. phymatum*, M9-fruc media was used. The results were processed in R program (https://www.r-project.org/) using *ad hoc* scripts.

In order to evaluate the termination efficiency at the single cell level, flow cytometry analysis was made for terminators T1, T7 and T9 in all hosts. For this purpose, a single colony from *P. putida, E. coli, Pseudomonas* UYIF39, *Pseudomonas* UYIF41 and *B. phymathum* strains harboring each of these constructs, plasmid pSEVA231 and plasmid pSEVA231-*Pj100*GFP were growth overnight in M9 media as aforementioned. Cells were diluted 1:10 in fresh M9 appropiate media and grown for an additional 6 hours. Cultures were stored in ice and sequentially analyzed on the Millipore Guava EasyCyte Mini Flow Cytometer, to quantify GFP fluorescence. For each sample three independent experiments were made. The data generated was processed with R scripts (https://www.r-project.org/), using the flowCore and flowViz packages, available on Bioconductor (https://bioconductor.org/).

### Analysis of terminator sequences

Putative terminators were sequenced using primer SEVA231F (5’-TCC GTA TGT TGC ATC ACC-3’). Sequences were trimmed from primers and subject to comparison with NCBI database using the BLAST algorithm (Blastn and Blastp). Sequences were aligned using ClustalW ^35^. Putative secondary structure of the terminator sequences at the RNA level were predicted with Mfold ^36^ using default parameters. For the prediction of transcriptional terminators, programs ARNold (http://rna.igmors.u-psud.fr/toolbox/arnold/) and FindTerm ^25^ were used with default parameters.

## Supporting information

## Acknowledgments

The authors are thankful to lab colleagues for insightful discussion about this manuscript.

