## Supplementary material for "Expanding the toolbox of broad host-range transcriptional terminators for Proteobacteria through metagenomics"

### Supporting Information

**Figure S1.** Efficiency of T1 terminator in *P. putida* KT2440. 2
**Figure S2.** Characterization of additional putative terminators in *P. putida* KT2440. 3
**Figure S3.** Growth profile of bacterial strains harboring the different constructs. 4
**Table S1.** Sequences of putative transcriptional terminators from eDNA identified in this study 5
**Table S2.** Description of the ORFs contained in the metagenomic inserts and their sequence similarities. 7
**Table S3.** Strains and plasmids used in this study. 16
**References** 18

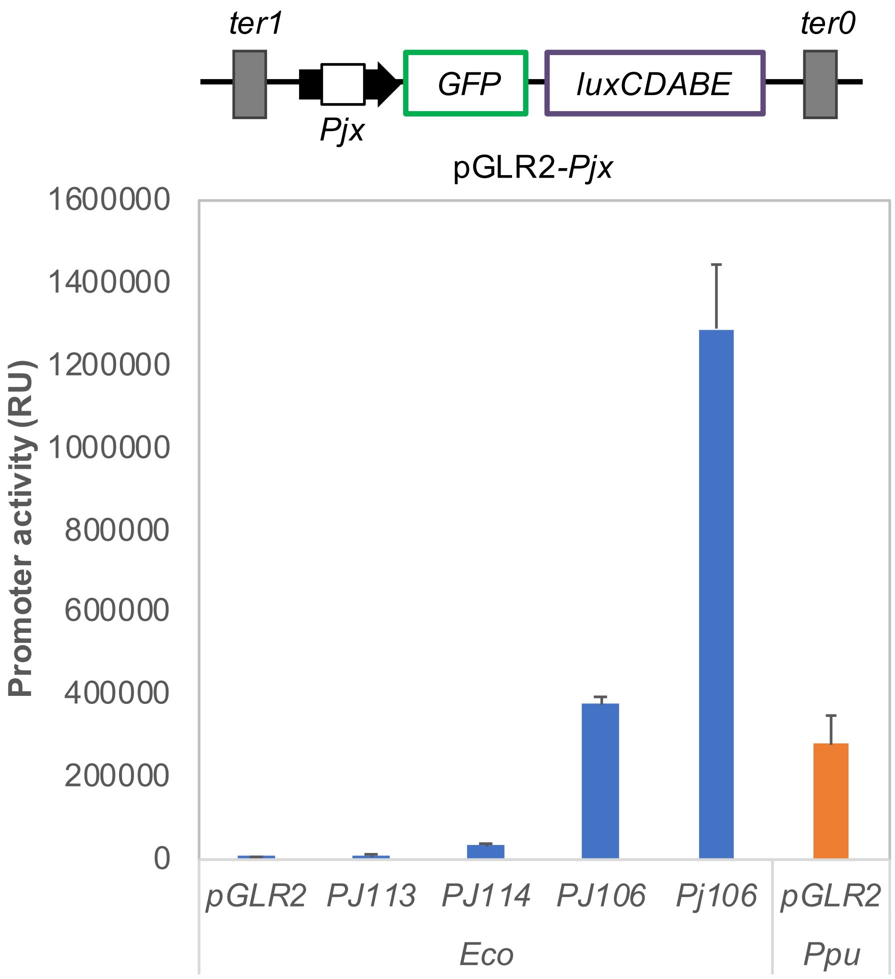


**Figure S1. Efficiency of T1 terminator in *P. putida KT2440*.** A reporter plasmid pGLR2 harboring different constitutive promoters were introduced in *E. coli* (*Eco*) and the promoter activity was assessed using the *lux* reporter [1]. The empty plasmid without promoter (pGLR2) was introduced in *P. putida* KT2440 (*Ppu*) for comparison. Activity observed in *P. putida* with the empty vector is as high as the promoter activity of *Pj106* variant in *E. coli*, indicating that the *ter1* terminator present in pGLR2 is not efficiently recognized in *P. putida*. Expression is putatively driven by plasmid intrinsic promoters driving the expression of other genes (i. e. antibiotic resistance gene). Values represent the average from three independent experiments. Vertical bars represent the standard deviations. RU: Relative units, calculated as luminescence normalized by $OD_{600}$.

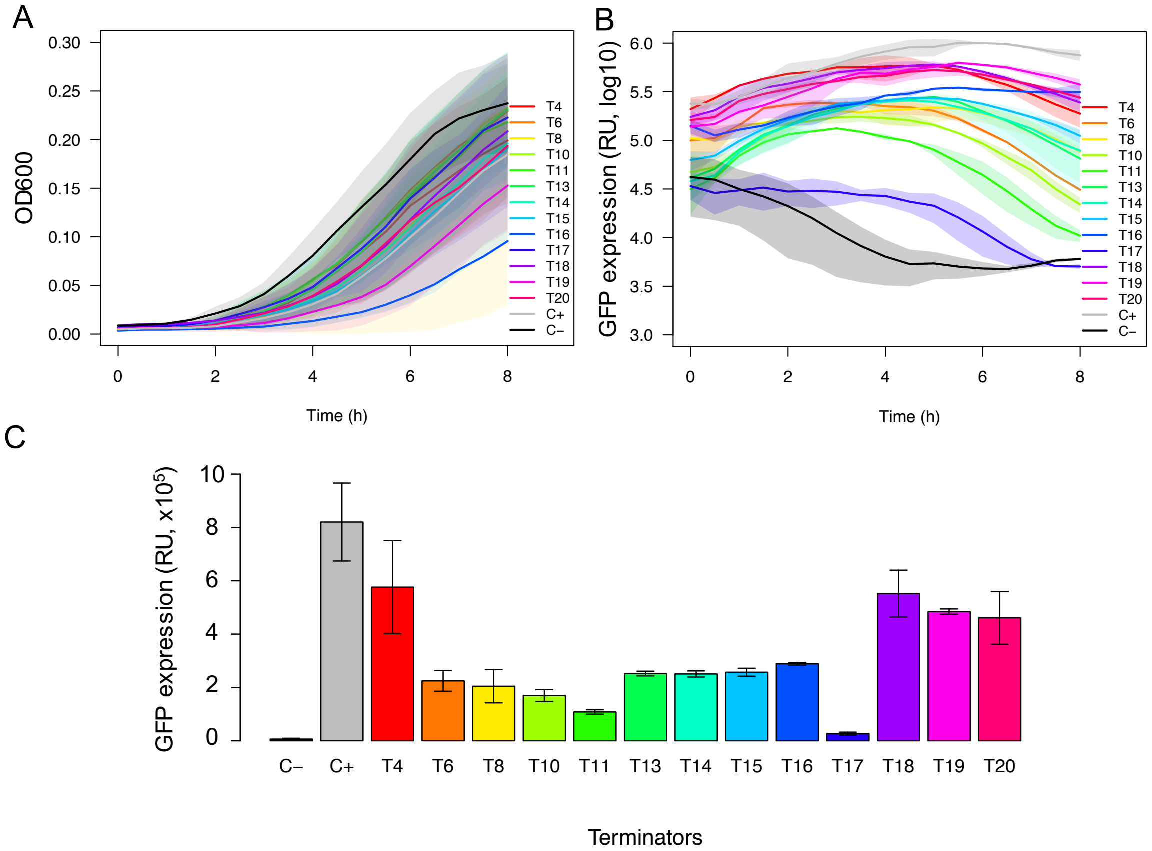


**Figure S2. Characterization of additional putative terminators in *P. putida* KT2440.** In order to characterize the transcriptional termination effect in the additional metagenomic clones found harboring putative transcriptional terminator sequences, *P. putida* KT2440 strains were grown in liquid M9-gly media during 8h and the optical density at 600 nm (OD) and fluorescence (GFP) were measured every 30 min. Strains harboring either pSEVA231 or pSEVA231-Pj100GFP were used as positive (C+) and negative (C-) controls, respectively. **A**) Growth curve of the strains; **B**) Terminator activity (in log10 scale) over time; **C**) Terminator activity at 4h. All graphs represent the average from three independent experiments. Standard deviation from three experiments is represented as shaded regions in A and B and as vertical bars C.

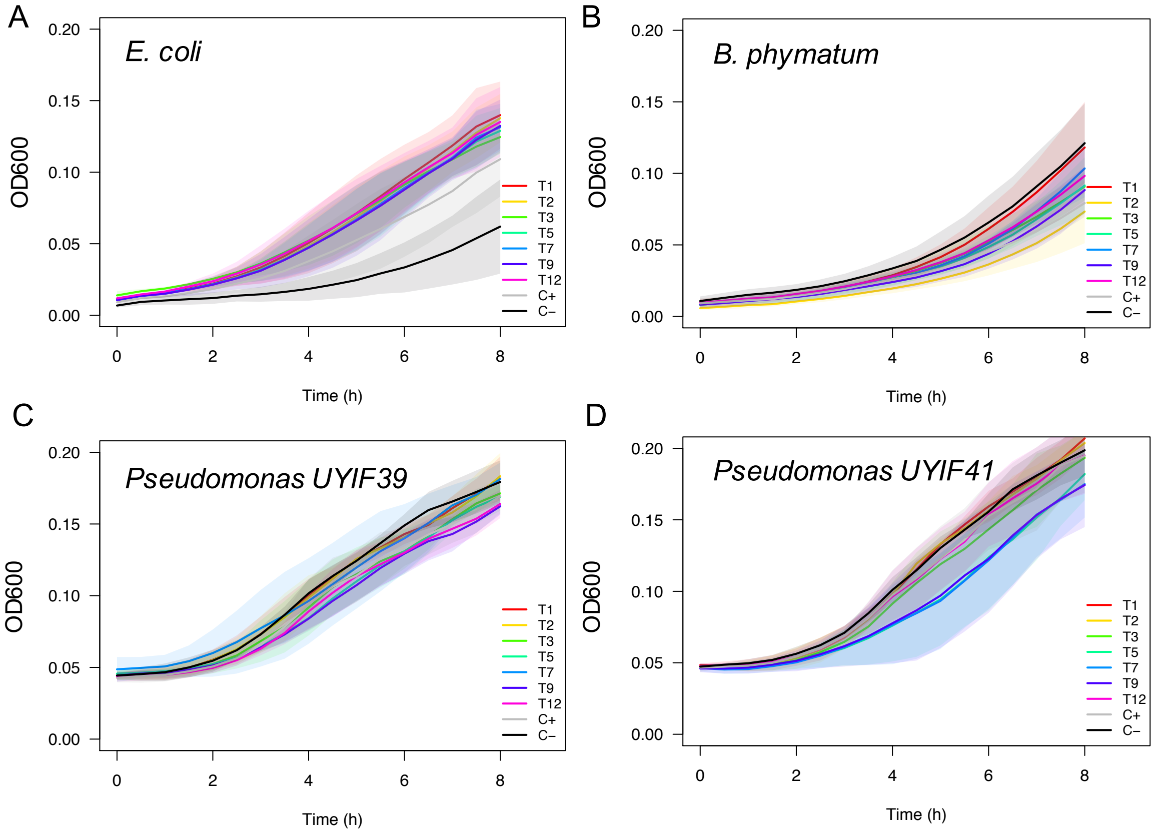


**Figure S3. Growth profile of bacterial strains harboring the** different constructs**.** Strains were grown in liquid M9 media supplemented with an appropriate carbon source during 8h and the optical density at 600 nm was measured every 30 min. Strains harboring either pSEVA231 or pSEVA231-Pj100GFP were used as positive (C+) and negative (C-) controls, respectively. All graphs represent the average from three independent experiments. **A)** Strains of *E. coli* DH10B. **B**) Strains of *Burkholderia phymatum* STM815$^{T}$. **C)** Strains of *Pseudomonas* UYIF39. **D)** Strains of *Pseudomonas* UYIF41. All graphs represent the average from three independent experiments.

**Table S1. Sequences of the putative transcriptional terminators from eDNA identified in this study**

| Name | Size (bp) | %GC | Sequence |
|---|---|---|---|
| T1 | 158 | 63.9 | 5´-<br>GAGCGCATGCTCGAGTACTTCGCCTGGACCATGCTCGCCGTCGTCTTCGGCTTCCTGCTCTTCGTCAACCTCGCCTACGTCCCCCTCGGCCA<br>CTGGGCCGAGACCTTCGCCGGCTTCTTCAAATTTTCAGGGCTGCCGCACCCCATCGACTGGGGGCT-3´ |
| T2 | 118 | 74.6 | 5´-<br>TCCGGGCCCGGTCACGGTCAGCGCGACGTCGTAGCTGCCCGCGGCGGCGTAGACGTGGGACGGGCTCGCGGACGTGGAGGTGCCGCCGT<br>CTCCGAACGTCCATGCCCAGGAGGCGACC-3´ |
| T3 | 355 | 68.2 | 5´-<br>TGCGCTTCGACGCGGTGCCCGAACGCGTCGACGTGCATATCATCAACCGTCCCGGCCAGCCCTTCCTCGGCAGCGGCGAAACCGGCCAGG<br>GCCCTGCGGCCGCTTCGATCGTGGACGGTCGGCTCCCTCGGGCGCCGATCTGCGATACGTTGAACTTTCAGCTGCTCGAGGTCGAGCGCG<br>GTCGCGCCGTATTTCAGGGCACGCCGAAGCCCGAGCACTACAACCCGCTCGGCTCGATCCATGGCGGTTGGCCCGCGACGCTGCTCGACT<br>CCTGCATGGCGTGCGCCGTGCACACCACGCTCGCCGCGGGCCAGGGCTATACGACGGTGGAGTTCAAGCTGAATCTGGTGCGGCC-3´ |
| T4 | 172 | 58.1 | 5´-<br>TGGGTTGGACCCGTTGCTTCGTGCAAGGTCGCCACGCTGTCCAGACCTATGCTCCCCAACCGCGTAGATGGAACGTTAAGACGCCGGATG<br>CTGGTTGCGTCGGTCCGGAAGTCCTCACTGGCACGAACGTGGACCTCATACTGCTCGCCGTTTTCGTTATAGGTCGAGACTT-3´ |
| T5 | 311 | 68 | 5´-<br>CCGTGGCGCAGCCTGCAAAACACAAAAAGCGCCCGGAGGAACGTCTCCTGAAACGCGTCTTCGGCCTGCTGCGGTCCGAGCCTCCGGCG<br>GAGGAGCGCCAGCACGATGCCTCGATGCTCCTCGTAGAAGGTCTCGAACGGCGGTATCGCCATCGCCTCCATCATGCTCTGAAACGGCCG<br>AGGCGGGCGCCGCGTGAGATCACCGCCCGCGCCGATCTCGATTCGCCGGTTCTTGGCGCGGCCCTCGGCGGTGTCATTCCCCAAGGGAGA<br>CCGGCTGTCGGCGAACGAGACCGCGGCCAGGCGCGTCTTGTC-3´ |
| T6 | 424 | 67.9 | 5´-<br>GTGCTCGATGGTCGAGAAGAGCAGAGGACGCCCGTCGTCAACCACCGTGGCGTTGAAGACTACGCCGGGACGGTTGCCCGCCCTTGCGTC<br>GCGCTTCCAGTCGCCGAGCATGGCTTTCAGATCCCGATGCCCGGCACCGTGCACGGCGCGTAGCCGCCGAACAGCGGCTGCGGAGTCTGG<br>AGCTTGCTGTTGTCGGCGATGCACATCAGCAGCACCATCGGCGCCTGGTGGGCGACCCAGGGCTGCACCGTCCCCACCCCCCCACGGTGC<br>TTGCCCGCGCCGCACGAGTCCGTCCAGTGCTGCGACAGCGGCACGAGGATCGTATGAAATTCGGCGACATCATCGCGCTCGCGTGGCGCA<br>ATCTCCGGCAGGCCAGGCTGAGAACCTCGTTGACGGTCATCGGCGTCGTCGTCGGCGTCGCGGC-3´ |
| T7 | 58 | 63.8 | 5´-CGCTCGTGCTCGCCGCGACCTTCCTCTTCATGCACCTCTTCGGCATCGGGCTGCACAA-3´ |
| T8 | 229 | 65.5 | 5´-<br>TCCGACCACGTTGTCGTCTCGTACCGCTCGCCGTGGACGTTCCTGACGTCGGCGTAGCGCGCGTAGAGCTGCGCGAGTGCGACGAAGCGA<br>TCGCCTCGTCATTGCGTCCATGGCCCGCCCACGCTTCGGCCATGTTCACGTAGGCCGAGACGATCCGCGGCGCATATTTGCGCGCACCGC<br>GGCGTCGCCGCTCTTGACCGCGTCGATCATCCAGGTCGAGTGCTTGAT-3´ |
| T9 | 181 | 54.7 | 5´-<br>GGAATCTCCTTCTCGCCTCTTTCGCTGAACACGAAAGCGAATCCGTTCAGCACACACTTTACGATATGGCGCAAAAAGTCCTCGGCTGCGT<br>GCCGGAAGTGAAAGACATCCACCTCACCATGCCCAACAAGCATTGCCTGCTCGTGGACCTGTCCCGCTTCGGTCAGGACAATCCCAACGA<br>-3´ |
| T10 | 175 | 54.9 | 5´- |

| | | | |
|---|---|---|---|
| | | | CCGTTTGATATTTGCCCCCGCTTTCAAAAAGAATGAGCTCGGCCCCAAGTTCGCGAATCAACATGCGCCGTTCCTCGCTGGCCGTAGCGGACATGAGTATTGTGACTGCATACCCTTTGGCGGCTCCCACCATTGAAAGCGCGATTCCGGTATTGCCGCTGGAGCATTCGAGGAT-3´ |
| T11 | 249 | 62.6 | 5´-TCCGCCACCTGCAGCAGGGCATGGACCACCGGCATCAACAACGGCGGAAAGCCGTCCACTCTTCCGCGAAGCCACACTTCCGGTTGCTCGGTCATAGCACCACAGTACCAGCGTCGGCCCGCCAAGCTCCAGATGTCTCTTACAATCGCGAGTGATGCCGCGAGACTACGTGCCGCAACTGGCGACGCTCGTCAAGACGCCACCGCAGGGCGACGGCTGGTGGCACGAAATCAAGTTCGACGGCTATCG-3´ |
| T12 | 129 | 69 | 5´-CTCGCTCTATCTCCATCACCCGACCGCGACCGACCCAAAAATGGCGCCGCCGGGCCATTCGACCTTCTACGCGCTCGCGCCGGTCCCCCATCTCGGCAAATTCCCCGTCGACTGGGCGCGCGTCGGGCC-3´ |
| T13 | 150 | 59.3 | 5´-TTCACCCGGCCAGGAGCGGTGTGCTAATTCCTCGTTGACCATCACCGCATCGGGCGCGTCGGCGCTGTCGCGTTCGGTAAACTCGCGCCCCTTTAGAAGCCGAATACCCATCGCACGAAAATAGCCAACGGTAATCGCGCGATAATCCGC-3´ |
| T14 | 251 | 60.2 | 5´-GTGGCAGCCGCCAAGGCCGCCTATGCCCACGATTTCATCATGTCCTTTCCGCGCGGCTATGACACACCGGTTGGCGAGCACGGCATGCAGCTCTCGGGTGGACAGCGTCAAAGAGTGGCTATCGCCCGCGCTCTCATTAAAAACGCACCGATCATTCTGCTTGACGAAGCAACGGCGGCGCTCGATTCCGAATCCGAGTTGCAGGTGCGTGAAGCCGTGGAACATCTCTGTCAAGGCCGCACGACGCTCGT-3´ |
| T15 | 372 | 56.9 | 5´-GGTGATTTCCTGGCGTACCAGTACGTCACTGACGTCAACTACAGCCGGGTGACCAACTTCAGCGAGATGGAGTTCGTCGTCCCCGGCCCCGGCGCCGTGGACGGGATACGTAAATGCTTTGCCGACACCGGGGGGCTGAACCACTCGGAAGTCATCCGGTTCATGGCCGACCGCCAAGAGATCGGTCTTTTGAGTCCGACCGCAGCGACATTTGCAACAAACTCCTTGAAGGCCTTTCGATTGCGGTCGCGAGGATTGACCACATCAATCAGATGCAAGGGGATTATCTCGATAGCAGGCAAGTTGGGAGGATCGCTCATGTAGCCTCCGCCAGTGTCGACCTCTCAATTAGGCTAATCACCCCCTCCAAGC-3´ |
| T16 | 208 | 57.2 | 5´-GCGACGAGAACTGCGACGGCATGCGCACCTGCTTCGCCAACGCCGACGGCGACATGTATCGGACGGCGATCAGGATGCTCGCCGCGCGAAATTCGGCAATTCCGCTTGCATGCAGGAGCCCAATTTGAAAAACGGCTGCGGCGGTCTGCGCGATTTTCAAAATCTGCTCTGGATGACTTTCTTCAAATATCGCACCCGCTCCCTCAAT-3´ |
| T17 | 152 | 50.7 | 5´-TGTTACCCAGCTCTGTAATTCGTGTGCAACCGAAGACATCCCTGACACTTCAATCCGAGGACATTCCTGACAGCTTTTGAGAGAGCTCTTTCTCCACCAGATCAGCCATCCTGCCTTTAGTTGGTGTGAAGGGGAGTGTAGGCACTTGTCG-3´ |
| T18 | 266 | 64.7 | 5´-ATCGCGCGCACCGCGGAAGAACTGCTGCTGCTCGTGCCGAACACGCTGCGCGAGGGAGCGTTGGCGCTGGGGGCCACGCGCGCGCGCGCGGTGTTCAGCGTGGTTCTTCCGGCAGCCGCACCAGGGATCGCTGCGTGCCGAGTGTACCGATGCACATCTGCGATGTCGTCTGCCGTACGAGGCTTGAAATCTGCCGCCGCTGGGTGGGAAGACACTACATCAATCAGCTAAAAAGGCCTCCGTTCGGGAAGAACGGAGGCCCAAGT-3´ |
| T19 | 224 | 68.7 | 5´-ACGTGGCGCTTGCCGGACGCGCCTTGCCGGAAGCGGCTCACGACGCCGTGAACGCTGCGCGCCATACCCCCCAAGCTGAGCGTGAGGGTCGCCGAGCGGCCGACGATATCGCGCAGCGCCGACGACGTCTCCTCGGTCGCCAGGGTCACCTGGAACTGAAACAGCTCCGAGATTGCCTCGACGCCCTCCAAGCGAACCAAGCGCAGCGCTTGGGAGACGCCGGT-3´ |
| T20 | 86 | 67.4 | 5´-TACGACCAGTGCAACTCGCTGCACACGAACGCCTACGACGAGGCGATCACGACGCCGACCGAGGAGTCGGTGCGCCGCGCGATGGC-3´ |

#### Table S2. Description of the ORFs contained in the metagenomic inserts and their sequence similarities

| Name | Size (bp) | %GC | Blastn (Accesion) | Query coverage (%) | Identity (%) | E-value | ORF | Strand | ORF sequence | Blastp (Accesion) | Query coverage (%) | Identity (%) | E-value |
|---|---|---|---|---|---|---|---|---|---|---|---|---|---|
| T1 | 158 | 63.9 | *Luteitalea pratensis* strain DSM 100886 (CP015136.1) | 100 | 82 | 4e-27 | 1 | + | MLEYFAWTMLAVVFGFLLFVNLAYVPLGHWAETFAGFFKFSGLPHPIDWG | hypothetical protein DMF83_08540 [*Acidobacteria bacterium*] (PYQ07790.1) | 100 | 82 | 1e-19 |
| | | | | | | | 2 | + | MDHARRRLRLPALRQPRLRPPRPLGRDLRRLLQIFRAAAPHRLGA | hypothetical protein [*Methylobacterium nodulans*] (WP_083786683.1) | 64 | 59 | 0.29 |
| T2 | 118 | 74.6 | NS[a] | NA | NA | NA | 1 | - | MGMDVRRRRHLHVREPVPRLRRRGQLRRRADRDRAR | NS | NA | NA | NA |
| T3 | 355 | 68.2 | *Bradyrhizobium erythrophlei* strain GAS138 (LT670817.1) | 31 | 88 | 3e-27 | 1 | + | MAVGPRRCSTPAWRAPCTPRSPRARAIRRWSSS | uncharacterized protein LOC105686051 isoform X2 [*Athalia rosae*] (XP_012256029.1) | 90 | 47 | 5.5 |
| | | | | | | | 2 | + | MRYVELSAARGRARSRRISGHAEARALQPARLDPWRLARDAARLLHGVRRAHHARRGPGLYDGGVQAESGAA | NS | NA | NA | NA |
| | | | | | | | 3 | + | MNFQLLEVERGRAVFQGTPKPEHYNPLGSIHGGWPATLLDSCMACAVHTTLAAGQGYTTVEFKLNLVR | PaaI family thioesterase [*Ideonella sakaiensis*] | 100 | 74 | 4e-29 |
| | | | | | | | 4 | - | MARGERGVHGARHAGVEQRRGPTAMDRAERVVVLGLRRALKYGATALDL | NS | NA | NA | NA |

| | | | | | | | | | | | | | |
|---|---|---|---|---|---|---|---|---|---|---|---|---|---|
| | | | | | | | | | E QLKVQRIADRRPREPTVHDRSGRRALAGFAAAEEGLAGTVDDMHVDAFGHRVEA | | | | |
| T4 | 172 | 58.1 | NS | NA | NA | NA | 1 | + | MSRPMLPNRVDGTLRRRMLVASVRKSSLARTWTSYCSPFSL | NS | NA | NA | NA |
| | | | | | | | 2 | + | MRRSGSPHWHERGPHTARRFRYRSRL | Serine/threonine-protein kinase pkn1 [*Acaryochloris sp.* RCC1774] (PZD74498.1) | 88 | 67 | 0.33 |
| | | | | | | | 3 | - | MRSTFVPVRTSGPTQPASGVLTFHLRGWGA | GAF domain-containing protein [*Streptomyces peucetius*] (WP_100110470.1) | 53 | 81 | 5.6 |
| T6 | 424 | 67.9 | *Micromonas commoda* predicted protein partial mRNA ( XM_002500349.1) | 6 | 100 | 0.009 | 1 | + | MLLSAMHISSTIGAWWATQGCTVPTPPRCLPAPHESVQCCDSGTRIV | hypothetical protein PV10_05849 [*Exophiala mesophila*] (XP_016222872.1) | 57 | 37 | 2.8 |
| | | | | | | | 2 | + | MVEKSRGRPSSTTVALKTTPGRLPALASRFQSPSMAFRSRCPAPCTARSRRTAAAESGACCCRRCTSAAPSAPGGRPRAAPSPPPHGACPRRTSPSSAATAARGSYEIRRHHRARVAQSPAGQAENLVDGHRRRRRRRG | NS | NA | NA | NA |
| | | | | | | | 3 | + | MRRASSRRAWLSDPDARHRARRVAAEQRLRSLEL | hypothetical protein DMG63_06990 | 29 | 61 | 0.001 |

| | | | | | | | | | | | | | |
|---|---|---|---|---|---|---|---|---|---|---|---|---|---|
| | | | | | | | | | AVVGDAHQQHHRRLVG<br>DPGLHRPHPPTVLARAA<br>RVRPVLRQRHEDRMKFG<br>DIIALAWRNLRQARLR<br>TSLTVIGVVVGVA | [*Acidobacteria bacterium*] (PYY00129.1) | | | |
| | | | | | | | 4 | - | MTVNEVLSLACRRLRHA<br>SAMMSPNFIRSSCRCRST<br>GRTRAARASTVGGWG<br>RCSPGSPTRRRWCC | NS | NA | NA | NA |
| | | | | | | | 5 | - | MDGLVRRGQAPWGGGD<br>GAALGRPPGADGAADV<br>HRRQQQAPDSAAAVRRL<br>RAVHGAGHRDLKAMLG<br>DWKRDARAGNRPGVVF<br>NATVVDDGRPLLFSTIEH | hypothetical protein [*Acaryochloris marina*] (WP_012168077.1) | 34 | 50 | 0.53 |
| | | | | | | | 6 | - | MPEIAPRERDDVAEFHTI<br>LVPLSQHWTDSCGAGKH<br>RGGVGTVQPWVAHQA<br>PMVLLMCIADNSKLQTP<br>QPLFGGYAPCTVPGIGI | hydantoinase B/oxoprolinase family protein [*Solirubrobacter soli*] (WP_028066685.1) | 79 | 97 | 1e-37 |
| T7 | 58 | 63.8 | NS | NA | NA | NA | 1 | - | LCSPMPKRCMKRKVAAS<br>TS | uncharacterized protein LOC106373213 [*Brassica napus*] (XP_013668878.1) | 94 | 58 | 21 |
| T8 | 229 | 65.5 | NS | NA | NA | NA | 1 | + | MARPRFGHVHVGRDDP<br>RRIFCAHRGVAALDRVD<br>HPGRVLD | NS | NA | NA | NA |
| | | | | | | | 2 | - | MIDAVKSGDAAVRAKY<br>APRIVSAYVNMAEAWA<br>GHGRNDEAIASSHSRSST<br>RATPTSGTSTASGTRRQR<br>GR | tetratricopeptide repeat protein [*Planctomycetes bacterium*] (REK10448.1) | 60 | 43 | 0.62 |
| | | | | | | | 3 | - | MDAMTRRSLRRTRAALR<br>ALRRRQERPRRAVRDDN<br>VVG | NS | NA | NA | NA |

| | | | | | | | | | | | | | |
|---|---|---|---|---|---|---|---|---|---|---|---|---|---|
| | | | | | | | 4 | - | MDDRRGQERRRRGARKICAADRLGLREHGRSVGGPWTQ | NS | NA | NA | NA |
| T9 | 181 | 54.7 | NS | NA | NA | NA | 1 | + | MNTKANPFSTHFTIWRKKSSAACRK | putative S-adenosyl-L-methionine-dependent methyltransferase [*Macleaya cordata*] (OVA14828.1) | 48 | 85 | 3.1 |
| | | | | | | | 2 | + | MRAGSERHPPHHAQQALPARGPVPLRSGQSQR | NS | NA | NA | NA |
| | | | | | | | 3 | + | MAQKVLGCVPEVKDIHLTMPNKHCLLVDLSRFGQDNPN | urate oxidase [*Acidobacteria bacterium*] (PYX20951.1) | 100 | 71 | 4e-11 |
| | | | | | | | 4 | - | MTEAGQVHEQAMLVGHGEVDVFHFRHAAEDFLRHIVKCVLNGFAFVFSERGEKEI | NS | NA | NA | NA |
| T10 | 175 | 54.9 | NS | NA | NA | NA | 1 | + | MIFAPAFKKNELGPKFANQHAPFLAGRSGHEYCDCIPFGGSHH | NS | NA | NA | NA |
| | | | | | | | 2 | + | MPPLSKRMSSAPSSRINMRRSSLAVADMSIVTAYPLAAPTIESAIPVLPLEHSR | Uncharacterised protein [*Chlamydia trachomatis*] (CRH77335.1) | 81 | 57 | 3e-05 |
| | | | | | | | 3 | - | MVGAAKGYAVTILMSATASEERRMLIRELGAELILFESGGKYQT | cysteine synthase A [*Verrucomicrobia bacterium*] (PYJ54962.1) | 100 | 77 | 4e-14 |
| | | | | | | | 4 | - | MLQRQYRNRAFNGGSRQRVCSHNTHVRYGQRGTAHVDSRTWGRAHSF | NS | NA | NA | NA |
| T11 | 249 | 62.6 | NS | NA | NA | NA | 1 | + | MSLTIASDAARLRAATGDARQDATAGRRLVARNQVRRLS | NS | NA | NA | NA |

| | | | | | | | | | | | | | |
|---|---|---|---|---|---|---|---|---|---|---|---|---|---|
| | | | | | | | 2 | + | MPRDYVPQLATLVKTPPQGDGWWHEIKFDGY | DNA ligase D [*Acidobacteria bacterium*] (PYR63411.1) | 90 | 82 | 4e-08 |
| | | | | | | | 3 | + | MQQGMDHRHQQRRKAVHSSAKPHFRLLGHSTTVPASARQAPDVSYNRE | NS | NA | NA | NA |
| | | | | | | | 4 | - | MELGGPTLVLWCYDRATGSVASRKSGRLSAVVDAGGPCPAAGGG | NS | NA | NA | NA |
| | | | | | | | 5 | - | MAGRRWYCGAMTEQPEVWLRGRVDGFPPLLMPVVHALLQVA | DinB family protein [*Algoriphagus yeomjeoni*] (WP_111610207.1) | 70 | 72 | 7e-06 |
| T12 | 129 | 69 | *Sphingomonas indica* strain Dd16 (LT840185.1) | 89 | 82 | 3e-17 | 1 | + | MAPPGHSTFYALAPVPHLGKFPVDWARVG | phytoene desaturase [*Sphingomonas mali*] (WP_066827803.1) | 89 | 96 | 1e-16 |
| | | | | | | | 2 | - | MGDRRERVEGRMARRRHFWVGRGRVMEIER | hypothetical protein DL98DRAFT_587228 [*Cadophora sp.* DSE1049] (PVH81613.1) | 80 | 57 | 0.009 |
| T13 | 150 | 59.3 | NS | NA | NA | NA | 1 | + | MTITASGASALSRSVNSRPFRSRIPIARK | uncharacterized protein Dpse_GA15140, isoform A [*Drosophila pseudoobscura pseudoobscura*] (XP_001354708.2) | 62 | 53 | 1.3 |
| | | | | | | | 2 | - | MGIRLLKGREFTERDSADAPDAVMVNEELAHRSWPGE | hypothetical protein DMG13_06910 [*Acidobacteria bacterium*] (PYS54766.1) | 100 | 76 | 4e-10 |

| | | | | | | | | | | | | |
|---|---|---|---|---|---|---|---|---|---|---|---|---|
| T14 | 251 | 60.2 | NS | NA | NA | NA | 1 | + | MSFPRGYDTPVGEHGMQLSGGQRQRVAIARALIKNAPIILLDEATAALDSESELQVREAVEHLCQGRTTL | ABC transporter ATP-binding protein [*Afipia broomeae*] (WP_006021537.1) | 100 | 89 | 2e-35 |
| | | | | | | | 2 | + | MPTISSCPFRAAMTHRLASTACSSRVDSVKEWLSPALSLKTHRSFCLTKQRRRSIPNPSCRCVKPWNISVKAARRS | NS | NA | NA | NA |
| | | | | | | | 3 | - | MTEMFHGFTHLQLGFGIERRRCFVKQNDRCVFNESAGDSHSLTLSTRELHAVLANRCVIAARKGHDEIVGIGGLGGCH | Uncharacterised protein [*Shigella sonnei*] (CSG17181.1) | 76 | 42 | 1e-07 |
| T15 | 372 | 56.9 | NS | NA | NA | NA | 1 | + | MAYQYVTDVNYSRVTNFSEMEFVVPGPGAVDGIRKCFADTGGLNHSEVIRFMADRQEIGLLSPTAATFATNSLKAFRLRSRGLTTSIRCKGIISIAGKLGGSLM | hypothetical protein [*Ponticaulis koreensis*] (WP_051134783.1) | 55 | 78 | 1e-26 |
| | | | | | | | 2 | + | MLCRHRGAEPLGSHPVHGRPPRDRSFESDRSDICNKLLEGLSIAVARIDHINQMQGDYLDSRQVGRIAHVASASVDLSIRLITPSK | NS | NA | NA | NA |
| | | | | | | | 3 | - | MRGRHWRRLHERSSQLACYRDNPLASD | PepSY domain-containing protein [*Sphingomonas panacis*] (WP_069206517.1) | 66 | 67 | 0.19 |
| | | | | | | | 4 | - | MAVGHEPDDFRVVQPPGVGKAFTYPVHGAGAGDDELHLAEVGHPAVVDVS | NS | NA | NA | NA |

| | | | | | | | | | | | | | |
|---|---|---|---|---|---|---|---|---|---|---|---|---|---|
| | | | | | | | | | DVLVRQEIT | | | | |
| | | | | | | | 5 | - | MSDPPNLPAIEIIPLHLIDVVNPRDRNRKAFKEFVANVAAVGLKRPISWRSAMNRMTSEWFSPPVSAKHLRIPSTAPGPGTTNSISLKLVTRL | chromosome partitioning protein ParB [*Bradyrhizobium erythrophlei*] (WP_079601058.1) | 46 | 63 | 1e-08 |
| | | | | | | | 6 | - | MMWSILATAIERPSRSLLQMSLRSDSKDRSLGGRP | NS | NA | NA | NA |
| T16 | 208 | 57.2 | NS | NA | NA | NA | 1 | + | MHAGAQFEKRLRRSARFSKSALDDFLQISHPLPQ | NS | NA | NA | NA |
| | | | | | | | 2 | + | MRRHAHLLRQRRRRHVSDGDQDARRAKFGNSACMQEPNLKNGCGGLRDFQNLLWMTFFKYRTRSLN | Bifunctional uridylyltransferase/uridylyl-removing enzyme [*Verrucomicrobia bacterium*] (SPE56715.1) | 69 | 91 | 5e-22 |
| | | | | | | | 3 | + | MRTCFANADGDMYRTAIRMLAARNSAIPLACRSPI | NS | NA | NA | NA |
| | | | | | | | 4 | - | MKIAQTAAAVFQIGLLHASGIAEFRAASILIAVRYMSPSALAKQVRMPSQFSS | ATP-dependent RNA helicase DbpA [*Sulfurimonas sp.* RIFOXYB12_FULL_35_9] (OHE03472.1) | 96 | 33 | 8 |
| T17 | 152 | 50.7 | NS | NA | NA | NA | 1 | + | MRELFLHQISHPAFSWCEGECRHLS | putative uncharacterized protein [*Ruminococcus sp.* CAG:254] (CCZ83550.1) | 76 | 61 | 12 |
| | | | | | | | 2 | - | MIWWRKSSLKSCQECPRIEVSGMSSVAHELQSWV | NS | NA | NA | NA |

| | | | | | | | | | | | | | |
|---|---|---|---|---|---|---|---|---|---|---|---|---|---|
| | | | | | | | | | T | | | | |
| T18 | 266 | 64.7 | NS | NA | NA | NA | 1 | + | MLLLVPNTLREGALALGATRARAVFSVVLPAAAPGIAACRVYRCTSAMSSAVRGLKSAAAGWEDTTSIS | phosphate ABC transporter, permease protein PstA [*Acidobacteria bacterium* 13_1_20CM_2_65_9] (OLE85201.1) | 52 | 89 | 3e-10 |
| | | | | | | | 2 | + | MPPLGGKTLHQSAKKASVREERRPK | hypothetical protein PHYSODRAFT_506723 [*Phytophthora sojae*] (XP_009528675.1) | 68 | 68 | 17 |
| | | | | | | | 3 | + | MHICDVVCRTRLEICRRWVGRHYINQLKRPPFGKNGGPS | NS | NA | NA | NA |
| | | | | | | | 4 | - | MIDVVSSHPAAADFKPRTADDIADVHRYTRHAAIPGAAAGRTTLNTARARVAPSANAPSRSVFGTSSSSSSAVRA | NS | NA | NA | NA |
| T19 | 224 | 68.7 | NS | NA | NA | NA | 1 | + | MRAIPPKLSVRVAERPTISRSADDVSSVARVTWN | NS | NA | NA | NA |
| | | | | | | | 2 | + | MPEAAHDAVNAARHTPQAEREGRRAADDIAQRRRRLLGRQGHLELKQLRDCLDALQANQAQRLGDAG | glycine/betaine ABC transporter substrate-binding protein [*Curtobacterium sp.* BH-2-1-1] (WP_083295151.1) | 49 | 48 | 6.8 |
| | | | | | | | 3 | - | MRLVRLEGVEAISELFQFQVTLATEETSSALRDIVGRSATLTLSLGGMARSVHGVVSRFRQGASGKRH | hypothetical protein BE21_02605 [*Sorangium cellulosum*] (KYG07338.1) | 98 | 51 | 4e-13 |
| T20 | 86 | 67.4 | *Azoarcus sp.* | 100 | 94 | 2e-27 | 1 | + | MHTNAYDEAITTPTEESV | methylmalonyl-CoA | 100 | 100 | 1e-14 |

| | | |
|---|---|---|
| KH32C (AP012304.1) | RRAM | mutase domain protein [*Leptospira interrogans* str. L1207] |

[a]NS: No significant similarity found
[b]NA: Not applicable

**Table S3.** Strains and plasmids used in this study.

| Strain or plasmid | Relevant characteristics | References |
|---|---|---|
| **Strains** | | |
| ***Pseudomonas*** | | |
| *P. putida* KT2440 | Reference strain. | [2] |
| KT2440 pSEVA231 | *P. putida* KT2440 harboring plasmid pSEVA231 ($Km^R$) | This work |
| KT2440 pSEVA231-*Pj100*GFP | *P. putida* KT2440 harboring plasmid pSEVA231- *Pj100*GFP ($Km^R$) | This work |
| KT2440 T1 | *P. putida* KT2440 harboring plasmid pSEVA231-*Pj100**T1*GFP ($Km^R$) | This work |
| KT2440 T2 | *P. putida* KT2440 harboring plasmid pSEVA231-*Pj100**T2*GFP ($Km^R$) | This work |
| KT2440 T3 | *P. putida* KT2440 harboring plasmid pSEVA231-*Pj100**T3*GFP ($Km^R$) | This work |
| KT2440 T4 | *P. putida* KT2440 harboring plasmid pSEVA231-*Pj100**T4*GFP ($Km^R$) | This work |
| KT2440 T5 | *P. putida* KT2440 harboring plasmid pSEVA231-*Pj100**T5*GFP ($Km^R$) | This work |
| KT2440 T6 | *P. putida* KT2440 harboring plasmid pSEVA231-*Pj100**T6*GFP ($Km^R$) | This work |
| KT2440 T7 | *P. putida* KT2440 harboring plasmid pSEVA231-*Pj100**T7*GFP ($Km^R$) | This work |
| KT2440 T8 | *P. putida* KT2440 harboring plasmid pSEVA231-*Pj100**T8*GFP ($Km^R$) | This work |
| KT2440 T9 | *P. putida* KT2440 harboring plasmid pSEVA231-*Pj100**T9*GFP ($Km^R$) | This work |
| KT2440 T10 | *P. putida* KT2440 harboring plasmid pSEVA231-*Pj100**T10*GFP ($Km^R$) | This work |
| KT2440 T11 | *P. putida* KT2440 harboring plasmid pSEVA231-*Pj100**T11*GFP ($Km^R$) | This work |
| KT2440 T12 | *P. putida* KT2440 harboring plasmid pSEVA231-*Pj100**T12*GFP ($Km^R$) | This work |
| KT2440 T13 | *P. putida* KT2440 harboring plasmid pSEVA231-*Pj100**T13*GFP ($Km^R$) | This work |
| KT2440 T14 | *P. putida* KT2440 harboring plasmid pSEVA231-*Pj100**T14*GFP ($Km^R$) | This work |
| KT2440 T15 | *P. putida* KT2440 harboring plasmid pSEVA231-*Pj100**T15*GFP ($Km^R$) | This work |
| KT2440 T16 | *P. putida* KT2440 harboring plasmid pSEVA231-*Pj100**T16*GFP ($Km^R$) | This work |
| KT2440 T17 | *P. putida* KT2440 harboring plasmid pSEVA231-*Pj100**T17*GFP ($Km^R$) | This work |
| KT2440 T18 | *P. putida* KT2440 harboring plasmid pSEVA231-*Pj100**T18*GFP ($Km^R$) | This work |
| KT2440 T19 | *P. putida* KT2440 harboring plasmid pSEVA231-*Pj100**T19*GFP ($Km^R$) | This work |
| KT2440 T20 | *P. putida* KT2440 harboring plasmid pSEVA231-*Pj100**T20*GFP ($Km^R$) | This work |
| *Pseudomonas sp.* UYIF39 | | [3] |
| UYIF39 pSEVA231 | *Pseudomonas* UYIF39 harboring plasmid pSEVA231 ($Km^R$) | This work |
| UYIF39 pSEVA231-*Pj100*GFP | *Pseudomonas* UYIF39 harboring plasmid pSEVA231-*Pj100*GFP ($Km^R$) | This work |
| UYIF39 T1 | *Pseudomonas* UYIF39 harboring plasmid pSEVA231-*Pj100**T1*GFP ($Km^R$) | This work |

| | | |
|---|---|---|
| UYIF39 T2 | *Pseudomonas* UYIF39 harboring plasmid pSEVA231-*Pj100**T2*GFP ($Km^R$) | This work |
| UYIF39 T3 | *Pseudomonas* UYIF39 harboring plasmid pSEVA231-*Pj100**T3*GFP ($Km^R$) | This work |
| UYIF39 T5 | *Pseudomonas* UYIF39 harboring plasmid pSEVA231-*Pj100**T5*GFP ($Km^R$) | This work |
| UYIF39 T7 | *Pseudomonas* UYIF39 harboring plasmid pSEVA231-*Pj100**T7*GFP ($Km^R$) | This work |
| UYIF39 T9 | *Pseudomonas* UYIF39 harboring plasmid pSEVA231-*Pj100**T9*GFP ($Km^R$) | This work |
| UYIF39 T12 | *Pseudomonas* UYIF39 harboring plasmid pSEVA231-*Pj100**T12*GFP ($Km^R$) | This work |
| *Pseudomonas sp.* UYIF41 | | [3] |
| UYIF41 pSEVA231 | *Pseudomonas* UYIF41 harboring plasmid pSEVA231 ($Km^R$) | This work |
| UYIF41 pSEVA231-*Pj100*GFP | *Pseudomonas* UYIF41 harboring plasmid pSEVA231-*Pj100*GFP ($Km^R$) | This work |
| UYIF41 T1 | *Pseudomonas* UYIF41 harboring plasmid pSEVA231-*Pj100**T1*GFP ($Km^R$) | This work |
| UYIF41 T2 | *Pseudomonas* UYIF41 harboring plasmid pSEVA231-*Pj100**T2*GFP ($Km^R$) | This work |
| UYIF41 T3 | *Pseudomonas* UYIF41 harboring plasmid pSEVA231-*Pj100**T3*GFP ($Km^R$) | This work |
| UYIF41 T5 | *Pseudomonas* UYIF41 harboring plasmid pSEVA231-*Pj100**T5*GFP ($Km^R$) | This work |
| UYIF41 T7 | *Pseudomonas* UYIF41 harboring plasmid pSEVA231-*Pj100**T7*GFP ($Km^R$) | This work |
| UYIF41 T9 | *Pseudomonas* UYIF41 harboring plasmid pSEVA231-*Pj100**T9*GFP ($Km^R$) | This work |
| UYIF41 T12 | *Pseudomonas* UYIF41 harboring plasmid pSEVA231-*Pj100**T12*GFP ($Km^R$) | This work |
| ***Escherichia coli*** | | |
| *E. coli* DH10B | mcrA Δmrr-hsdRMS-mcrBC) φ 80lacZΔM15 ΔlacX74 recA1 araD139 Δ(ara-leu)7697 galU galK rpsL endA1 nupG Δdcm. | [4] |
| DH10B pMR1-*Pj100* | *E. coli* DH10B harboring plasmid pMR1-*Pj100* | [1] |
| DH10B pSEVA231 | *E. coli* DH10B harboring plasmid pSEVA231 ($Km^R$) | This work |
| DH10B pSEVA231-*Pj100*GFP | *E. coli* DH10B harboring plasmid pSEVA231-*Pj100*GFP ($Km^R$) | This work |
| DH10B T1 | *E. coli* DH10B harboring plasmid pSEVA231-*Pj100**T20*GFP ($Km^R$) | This work |
| DH10B T2 | *E. coli* DH10B harboring plasmid pSEVA231-*Pj100**T20*GFP ($Km^R$) | This work |
| DH10B T3 | *E. coli* DH10B harboring plasmid pSEVA231-*Pj100**T20*GFP ($Km^R$) | This work |
| DH10B T5 | *E. coli* DH10B harboring plasmid pSEVA231-*Pj100**T20*GFP ($Km^R$) | This work |
| DH10B T7 | *E. coli* DH10B harboring plasmid pSEVA231-*Pj100**T20*GFP ($Km^R$) | This work |
| DH10B T9 | *E. coli* DH10B harboring plasmid pSEVA231-*Pj100**T20*GFP ($Km^R$) | This work |
| DH10B T12 | *E. coli* DH10B harboring plasmid pSEVA231-*Pj100**T20*GFP ($Km^R$) | This work |
| ***Burkholderia*** | | |
| *Burkholderia* | Type strain. Accession number: PRJNA17409 | [5] |

| | | |
|---|---|---|
| *phymatum* STM815$^T$ | | |
| STM815 pSEVA231 | *B. phymatum* STM815 harboring plasmid pSEVA231 (Km$^R$) | This work |
| STM815 pSEVA231-*Pj100*GFP | *B. phymatum* STM815 harboring plasmid pSEVA231-*Pj100*GFP (Km$^R$) | This work |
| STM815 T1 | *B. phymatum* STM815 harboring plasmid pSEVA231-*Pj100**T1*GFP (Km$^R$) | This work |
| STM815 T2 | *B. phymatum* STM815 harboring plasmid pSEVA231-*Pj100**T2*GFP (Km$^R$) | This work |
| STM815 T3 | *B. phymatum* STM815 harboring plasmid pSEVA231-*Pj100**T3*GFP (Km$^R$) | This work |
| STM815 T5 | *B. phymatum* STM815 harboring plasmid pSEVA231-*Pj100**T5*GFP (Km$^R$) | This work |
| STM815 T7 | *B. phymatum* STM815 harboring plasmid pSEVA231-*Pj100**T7*GFP (Km$^R$) | This work |
| STM815 T9 | *B. phymatum* STM815 harboring plasmid pSEVA231-*Pj100**T9*GFP (Km$^R$) | This work |
| STM815 T12 | *B. phymatum* STM815 harboring plasmid pSEVA231-*Pj100**T12*GFP (Km$^R$) | This work |
| **Plasmids** | | |
| pMR1-*Pj100* | pMR1 harboring the strong synthetic constitutive promoter BBa_J23100 (*Pj100*) cloned as an EcoRI/BamHI fragment (Cm$^R$) | [1] |
| pSEVA231 | Cloning vector (Km$^R$). Accession number: JX560328 | [6] |
| pSEVA231-*Pj100*GFP | Terminator trap vector. pSEVA231 harboring a 840 bp DNA sequence form pMR1-*Pj100* containing *gfplva* under the control of *Pj100* promoter (*Pj100*GFP) cloned as an EcoRI/HindIII fragment (Km$^R$) | This work |
| pSEVA231-*Pj100*Tx**GFP | Terminator trap vector pSEVA231- *Pj100*GFP with eDNA sequences cloned at the BamHI site. Tx corresponds to different eDNA sequences with terminator function, where x is an arbitrary number from 1 to 20. | This work |

(5) Moulin, L.; Klonowska, A.; Caroline, B.; Booth, K.; Vriezen, J. a C.; Melkonian, R.; James, E. K.; Young, J. P. W.; Bena, G.; Hauser, L.; et al. Complete Genome

Sequence of Burkholderia Phymatum STM815(T), a Broad Host Range and Efficient Nitrogen-Fixing Symbiont of Mimosa Species. *Stand. Genomic Sci.* **2014**, *9* (3), 763–774. https://doi.org/10.4056/sigs.4861021.
(6) Silva-Rocha, R.; Martínez-García, E.; Calles, B.; Chavarría, M.; Arce-Rodríguez, A.; de las Heras, A.; Páez-Espino, A. D.; Durante-Rodríguez, G.; Kim, J.; Nikel, P. I. The Standard European Vector Architecture (SEVA): A Coherent Platform for the Analysis and Deployment of Complex Prokaryotic Phenotypes. *Nucleic Acids Res.* **2013**, *41* (D1), D666–D675.
